# Rapid High-Throughput Method for Investigating Physiological Regulation of Neutrophil Extracellular Trap Formation

**DOI:** 10.1101/2023.12.12.571175

**Authors:** Kieran Zukas, Justin Cayford, Finley Serneo, Brandi Atteberry, Andrew Retter, Mark Eccleston, Theresa K Kelly

## Abstract

Neutrophils, the most abundant white blood cells in humans, play pivotal roles in innate immunity, rapidly migrating to sites of infection and inflammation to phagocytose, neutralize, and eliminate invading pathogens. Neutrophil Extracellular Trap (NET) formation in response to pathogens is increasingly recognized as an essential rapid innate immune response, but when dysregulated contributes to pathogenesis of sepsis and immunothrombotic disease. Current models of NETosis are limited, routinely employing non-physiological triggers that can bypass natural NET regulatory pathways. Models utilizing isolated neutrophils and immortalized cell lines, do not reflect the complex biology underlying neutrophil activation and NETosis, that occurs in whole-blood. Here we describe a novel, high-throughput *ex-vivo* whole blood induced NETosis model using combinatorial pooling of native NETosis inducing factors in a more biologically relevant Synthetic-Sepsis™ model. We found different combinations of factors evoked distinct neutrophil responses in the rate of NET generation and/or magnitude of NETosis. Despite inter-donor variability, similar sets of pro-inflammatory molecules induced consistent responses across donors. We found at least three biological triggers were necessary to induce NETosis in our system including either TNF-α or LT-α. To our knowledge, we report the first human *ex-vivo* model utilizing naturally occurring molecules to induce NETosis in whole blood. This approach could be used for drug screening and, importantly, inadvertent activators of NETosis. These findings emphasize the importance of investigating neutrophil physiology in a biologically relevant context to enable a better understanding of disease pathology, risk factors, and therapeutic targets, potentially, providing novel strategies for disease intervention and treatment.

**Essentials:**

- NETosis is a vital immune response, but dysregulation leads to disastrous health outcomes.
- Current NETosis models don’t reflect the complex endogenous signaling that occurs in whole blood
- Netosis induction stimuli differs between isolated neutrophils and whole blood.
- A minimum of three physiological factors are required to induce NETosis in whole blood.

## Background

The innate immune system is the body’s first line of defense and rapidly responds to infection. Neutrophils represent 50-70% of white blood cells in humans and play a pivotal role in the initial response to infection. Whilst neutrophils can phagocytose pathogens, they also release extracellular traps to rapidly immobilize pathogens and prevent dissemination. NETosis, first described by Brinkmann et al in 2004 [1], involves the formation of Neutrophil Extracellular Traps (NETs) through rapid decondensation of nuclear chromatin, driven by changes to histone post translational modifications, followed by externalization of web like NETs containing long strands of chromatin and include associated antimicrobial granular enzymes (neutrophil elastase (NE) and myeloperoxidase (MPO)) [1]. While NETs serve as a vital defense mechanism, mounting evidence suggests that dysregulation and excessive formation can contribute to pathogenesis in sepsis and other immunothrombotic disorders through host directed bystander effects, initiation of a hyperinflammatory feedback loop and Disseminated Intravascular Coagulation [2]. Elevated nucleosome levels, a component of NETs, have been described in several studies of NETosis related conditions including COVID-19 and sepsis, and are negatively correlated with survival [3-6].

NETosis research has largely relied on mouse models, *in-vitro* models using isolated primary neutrophils or neutrophil-like cells induced from immortalized cell lines [7-9]. While these models have provided valuable insights into NETosis, they have significant limitations. Murine immune responses, though informative, can differ substantially from human responses in part due to the significantly lower proportion of neutrophils, their maturation profile and half-life [10-12]. Immortalized HL-60 cells are highly dependent on culture conditions and do not recapitulate neutrophil fragility. NETosis induction in isolated primary neutrophils overcomes many of these limitations providing important insights into neutrophil activation and regulation [13-15]. However, neutrophils are very fragile and the method of isolation impacts their response to various stimuli [16]. To bridge the gap between existing models and the clinical reality of human immunothrombotic disease, relevant *ex vivo* human models that enable rapid processing and require minimal handling are essential [12].

Various synthetic, as well as physiologically relevant factors induce NETosis. *In-vivo*, Staphylococcus aureus and Lipopolysaccharides (LPS) are commonly used NETosis inducers, while *in-vitro* models typically use LPS or Calcium Ionophore (CI) and, the most commonly reported inducer, phorbol 12-myristate 13-acetate (PMA) [17]. PMA is an extremely powerful inducer of NETosis but is not physiologically relevant as it can bypass natural regulatory pathways governing NET production, thus preventing the ability to understand full regulatory feedback loops, limiting the clinical relevance of findings from PMA induced studies [18].

Despite advances in critical care, sepsis and immunothrombotic disorders remain major global health burdens. New strategies are urgently needed to unravel the intricacies of the pathophysiologies, to understand individual patient susceptibility and ultimately develop more effective diagnostic tools and treatments. An *ex-vivo* human NETosis model using physiologically relevant triggers in the presence of other blood cells (e.g. macrophages, platelets) and circulating proteins offers several distinct advantages. Primarily, investigation of the dynamics of NET formation, regulation and function in a clinical context would better translate insights to patient settings. Additionally, human cell models enable exploration of patient-specific factors including genetic predisposition and influence of pre-existing conditions, which can significantly impact sepsis and immunothrombotic outcomes. By dissecting the molecular mechanisms underlying NETosis using human cells, we can potentially identify novel therapeutic targets and develop personalized sepsis treatment strategies.

### Objectives

Here, we developed a human primary cell-based Synthetic-Sepsis™ model using intact whole blood to study NETosis induction with physiologically relevant molecules (Supplemental Figure 1A). We show NETosis induction using panels of pro-inflammatory molecules varied in both time-scale and magnitude of NET release compared to non-physiological PMA induction. Furthermore, we show differential NETosis profiles based on specific combinations of molecules, which we hypothesize could distinguish between beneficial and pathogenic NETosis. Moreover, we show that TNF-α or LT-α was necessary but not sufficient for NETosis induction and that in the presence of C5a, rapid onset of NETosis occurred within two hours of exposure. A minimal combination of LT-α, C5a and fMLP was able to consistently induce NETosis in multiple donors. We believe our novel model could delineate underlying complexities of NETosis, potentially leading to the development of innovative diagnostic tools and targeted interventions for immunothrombotic disorders and other NETosis related pathologies.

## Methods

### Whole Blood Acquisition

Anonymous healthy donor K2-EDTA whole blood was obtained from PrecisionMed (San Diego). Research was approved under WCG IRB Protocol number 20181025 and all human participants gave written informed consent. Subjects were self-declared healthy between the ages of 18-50 with BMI < 30 and not taking NSAIDs. Whole Blood was stored at room temperature (RT) and processed within one-hour post-draw.

### Neutrophil isolation & Imaging

Neutrophils were isolated from whole blood using the MACSxpress Whole Blood Neutrophil Isolation Kit (Miltenyi, 130-104-434), with erythrocyte lysis conducted using 0.22x PBS hypotonic lysis buffer, and 1.78x PBS equilibration buffer and the EasySep Direct Human Neutrophil Isolation kit (StemCell Tech #19666). The kits were used as suggested by the manufacturer. Neutrophil purity was confirmed by FACS (Supplementary figure 1B-C).

### FACS

Whole blood fixation and neutrophil isolation were performed following a standardized protocol. Briefly, whole blood samples were collected in tubes containing K2 EDTA. Fixation was achieved by adding a 10x formaldehyde solution or a 4% paraformaldehyde solution to the whole blood samples for 10 minutes. After incubation and quenching of fixation, cells were resuspended in ice-cold 1x PBS for further processing.

### Isolated Neutrophil NETosis Induction

Neutrophils were resuspended in RPMI (Gibco, 11-875-119) containing 250nM Cytotox Green (Sartorious, 4633) to 2.0 x 10^5 cells/mL and seeded at 100µL/well in a 96-well Incucyte ImageLock plates (Sartrious, 4806) coated with 10µg/mL Fibronectin (Sigma-Aldrich, F1141). The plate was centrifuged at 120*g* for two minutes to seat the neutrophils at the bottom of the plate. NETosis stimuli, PMA (Sigma Aldrich, P1585), CI (Sigma-Aldrich, C7522), Lipopolysaccharide from *Pseudomonas aeruginosa* 10 (LPS, Sigma-Aldrich, L7018), and inhibitors were diluted in RPMI containing Cytotox Green then added to the plate containing neutrophils. Inhibitors, 4-aminobenzoic acid hydrazide (ABAH, Sigma-Aldrich, A41909-10G), or diphenyleneiodonium chloride (DPI, Sigma-Aldrich, D2926-10MG), were incubated with neutrophils at 37°C, 5% CO_2_ for 30 minutes before adding stimuli. The plate was imaged every 20 minutes with an Incucyte S3 Live-Cell Analysis System (Sartorious) using the phase contrast and green fluorescent channels at a 10× objective lens. NETosis was analyzed by excluding objects smaller than 30 µm^2^ in the phase channel and measuring the total area of the green signal with Top-Hat for background correction and Edge Split off.

### NETosis Induction in Whole Blood

25mL whole blood was reoxygenated by tube rolling at room temperature in 50mL tubes.

Periodically, oxygen saturation levels were determined by deoxyhemoglobin (660nm) and oxyhemoglobin (940nm) absorbance measurements. Oxygenated whole blood was then treated with PMA, CI, LPS, physiological molecules (10µg/mL LT-α, 10µg/mL C5a, 10µg/mL fMLP) or vehicle(PBS or DMSO) followed by inversion mixing. 2mL of treated sample was aliquoted into low binding tubes (ThermoScientific, 90410) and incubated at 37°C with rotation at 10 RPM. Plasma was isolated by centrifugation (swinging bucket) for 10 minutes at 1300*g* without brake at RT. Plasma fractions were transferred isolated and nucleosomes measured in duplicate using the H3.1 Nu.Q^®^ NETs immunoassay (Belgian Volition).

### Screening

Whole blood screening candidates were selected based on possible association with NETosis (Supplementary Table 1). Concentrations were equivalent to or in excess of values commonly used for *in-vitro* stimulation. Recombinant lyophilized proteins were resuspended using 0.1% (w/v) human serum albumin (HSA) (A9731, Sigma-Aldrich) in sterile water, with further dilutions of reagents made using 0.1% HSA in PBS. Reagents were dispensed in plates (Corning, 3575) using the Mantis V3 Liquid Handler (Formulatrix) with Silicon LV or PFE LV chips (233581 or 233129, Formulatrix). Wells were backfilled with appropriate solvents (Ethanol, DMSO, 0.1% HSA in PBS), such that the concentration and volume of vehicle was consistent across wells.

Plates contained 6 wells of each of the following controls: untreated, PMA (50, 250, and 500 nM) randomly distributed across the plate to assess assay performance. The factors contained within remaining wells were determined by the optimal DOE screening design computed using JMP V17.1 (JMP Statistical Discovery).

### Fluorescent Plate Assay

50 µL of oxygenated whole blood containing 7.5µM Sytox Green (Thermo Scientific, S7020) was dispensed across a prepared 384-well fluorescent assay microplate using the Mantis Continuous Flow Silicon Chip (Formulatrix, 233127). After dispensing, the plate was sealed (ThermoScientific, 235307) and mixed (orbital shaking) at 1400RPM for 10 seconds. Plates were then centrifuged at 1300*g* for 10 minutes at RT using a swinging bucket rotor, without braking. Centrifuged plates were pre-heated to 37°C on a dry-heat block (Lonza, 25-038A) for 15 minutes prior to reading. Kinetic fluorescent measurements were obtained using a SpectraMax iD5 Multi-Mode MicroPlate Reader (Excitation = 510nm, Emission = 550nm), top read, at two-minute intervals for up to 24 hours.

For inhibition studies, ABAH, MeOSuc-AAPA-CMK (Elastase Inhibitor II, Sigma Aldrich, 324755), Cl, GSK484 (Sigma-Aldrich, SML-1658), DPI, Necrostatin-1 (Sigma-Aldrich, 480065), Caspase-3/7 Inhibitor I (Cayman Chemical, 14464), or vehicle were added to the wells. Oxygenated whole blood was added to each well, mixed at 1400RPM for 5 seconds followed by RT incubation for 45-60 minutes. Vehicle, PMA, or physiological molecules(10µg/mL LT-α (Peprotech, 300-01B), 10µg/mL C5a (Peprotech, 300-70), 10µg/mL N-Formyl-Met-Leu-Phe (fMLP, Sigma-Aldrich, F3506) were then dispensed into each well and mixed at 1400RPM for 5 seconds. The plate was sealed, and assay continued as described above.

### MPO-DNA assay

MPO-DNA complexes were measured from isolated plasma according to Pieterse et al [19].

### Screening Design and Statistical Analysis

Optimal screening design was computed using JMP V17.1 (JMP Statistical Discovery) custom screening design, with all main effects and fourth degree interactions included, but assigned estimatibility to “if possible”. The number of runs specified for each plate was set to 360 with no additional center points or replicate runs. RFU signal was down sampled into 30min intervals by block-wise averaging, followed by calculating the change in the down sampled RFU values between 30-minute intervals. These values were used for input into standard least squares multivariate regression modeling using JMP V17.1, with the concentration of each molecule in each treatment being the dependent variable, and each change in RFU value at 30-minute intervals being the independent variable. The effect of each factor was assessed by t-test to determine if the predicted coefficient at any given period was non-zero, and if the factor was acting to increase or decrease signal at any given time. Factors with a significant non-zero effect on the signal and acted to increase signal over time (P < 0.1) in one or both tested donors for that screen, were selected to move forward for additional screening.

Dose-Response profiles for the three-factor combination were characterized by space-filling DOE (JMP V17.1). The selected design was sphere-packing optimal with 88 different concentration combinations of LT-α, C5a, and fMLP and tested in triplicate. The system’s dose-response surface was generated using gaussian process regression and gaussian correlation structure, with nugget parameter estimation. Each individual factor’s EC_50_ was obtained by determining the log concentration at the signal half-max for each factor, given that the two other factors are at their maximal concentrations.

Assessment of inhibition for the biological factors was conducted by subtracting the mean RFU value for biological factors and inhibitor treatment, with the mean RFU value for its respective inhibitor treatment using 30-minute block average at 6 hours into reading the plate. The standard deviation of both the biological factors with inhibitor treatment and inhibitor treatment alone are propagated to the ΔRFU value obtained. Comparison of the response of the biological factors between the inhibitor vehicle treatment and inhibitor treatments was performed using One-way ANOVA followed by Dunnett’s multiple comparisons test using GraphPad Prism (version 10.1).

## Results

### NETosis Induction and real-time monitoring in whole blood and isolated neutrophils

We set out to develop a more biologically relevant NETosis model than isolated neutrophils. Our approach allows the investigation of NETosis neutrophils in whole blood either in a low throughput, Synthetic Sepsis™ or high-throughput screening approach (Supplementary **Figure 1A**). We first validated our NET quantification approach, using extracellular DNA intercalation and fluorescence, in a classical isolated neutrophil model of NETosis. A dose dependent increase in DNA release measured by Cytotox green was seen in response to PMA using an Incucyte S3 incubated imaging platform (Sartorius), with signal onset after 2 hours of treatment (**Figure 1A**). Treatment with the MPO inhibitor, ABAH, substantially delayed the onset of PMA induced DNA release and reduced the overall level of release, consistent with NETosis inhibition (**Figure 1B**).

**Figure 1.**
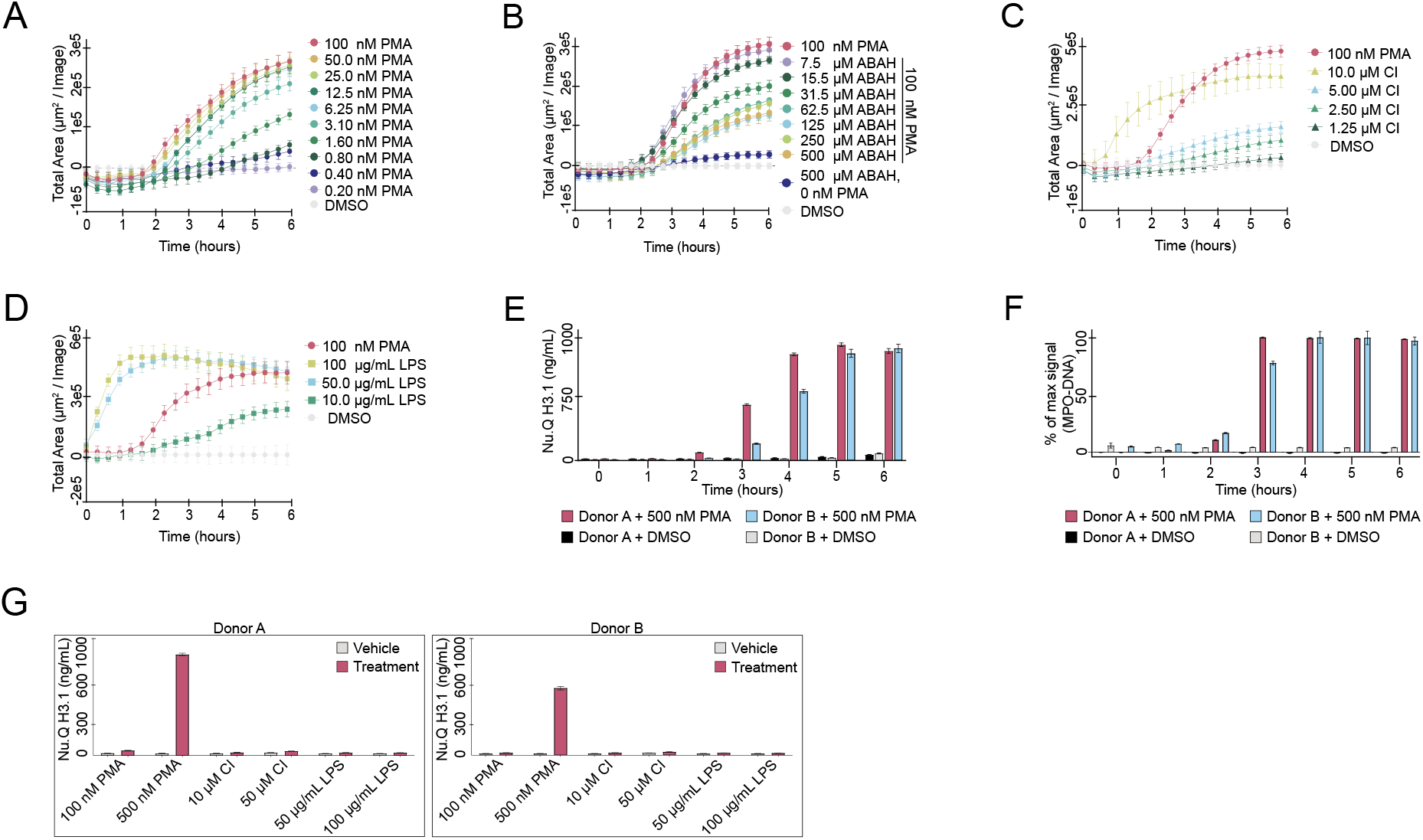
NETosis induction in isolated Neutrophils and whole-blood. (A) Phorbol myristate acetate (PMA) titration and DMSO vehicle control on isolated neutrophils using Cytotox green dye and S3 Incucyte imaging system. (B) Neutrophils were incubated with various concentrations of the myeloperoxidase inhibitor (ABAH) before the addition of 100 nM PMA. Controls included vehicle (DMSO) only, 100 nM PMA only and 500 µM ABAH only. (C) Isolated neutrophils were treated with various concentrations of Calcium Ionophore (CI) (▴) and compared to a DMSO control (grey ·) and 100 nM PMA (red ·). (D) Isolated neutrophils were treated with various concentrations of lipopolysaccharide (LPS) (◼) and compared to a DMSO vehicle control (grey ·) and 100 nM PMA (red ·). (E) Plasma nucleosome levels were measured using H3.1 Nu.Q® following 500 nM PMA addition to whole blood treated over a 0-6 hour time course. Donor A is indicated by red (treatment) or black (DMSO control). Donor B is indicated by blue (treatment) or grey (DMSO control). (F) MPO-DNA levels were measured by ELISA in same plasma samples as 1E. Bars are represented as percentage of max OD signal. (G) Whole blood from two donors were treated with low or high doses of PMA, CI, or LPS and H3.1 Nu.Q® levels were measured after 4 hours. Left panel is Donor A and the right panel is Donor B.

The results were confirmed by fluorescent microscopy at 6 hours post treatment (Supplementary **Figure 1E**). CI and LPS, established NETosis inducers, also induced NETosis in isolated neutrophils in a dose dependent manner (**Figure 1C and Figure 1D**).

To study NETs in a more biologically relevant system, (i.e. in the presence of other blood proteins and cell types) we incubated whole blood in K2-EDTA tubes from two healthy donors with PMA, CI or LPS. NETosis was quantified by nucleosome release and H3.1 nucleosome levels started to increase three hours after treatment with PMA (**Figure 1E**) as did MPO-DNA (**Figure 1F**). Differential nucleosome elevation was seen in both donors over the time course. Unlike in isolated neutrophils, where all three molecules induced NETosis, only PMA induced NETosis in whole blood in either donor (**Figure 1G**). This unexpected finding suggests that neutrophils respond differently in isolation than in whole blood.

### High-throughput model for NETosis induction

To identify physiologically relevant NETosis activators we adapted the model for high-throughput screening. As neutrophils are fragile, we designed a rapid screening format with limited handling utilizing a MANTIS liquid dispenser (Formulatrix) to distribute molecules across a 384-well plate followed by whole blood containing sytox green. The plate was centrifuged to sediment erythrocytes leaving an upper plasma layer and buffy coat containing neutrophils at the interface (**Figure 2A**). NET formation was measured from the top through the plasma via intercalation and fluorescence of extracellular DNA using a SpectraMax plate reader.

**Figure 2.**
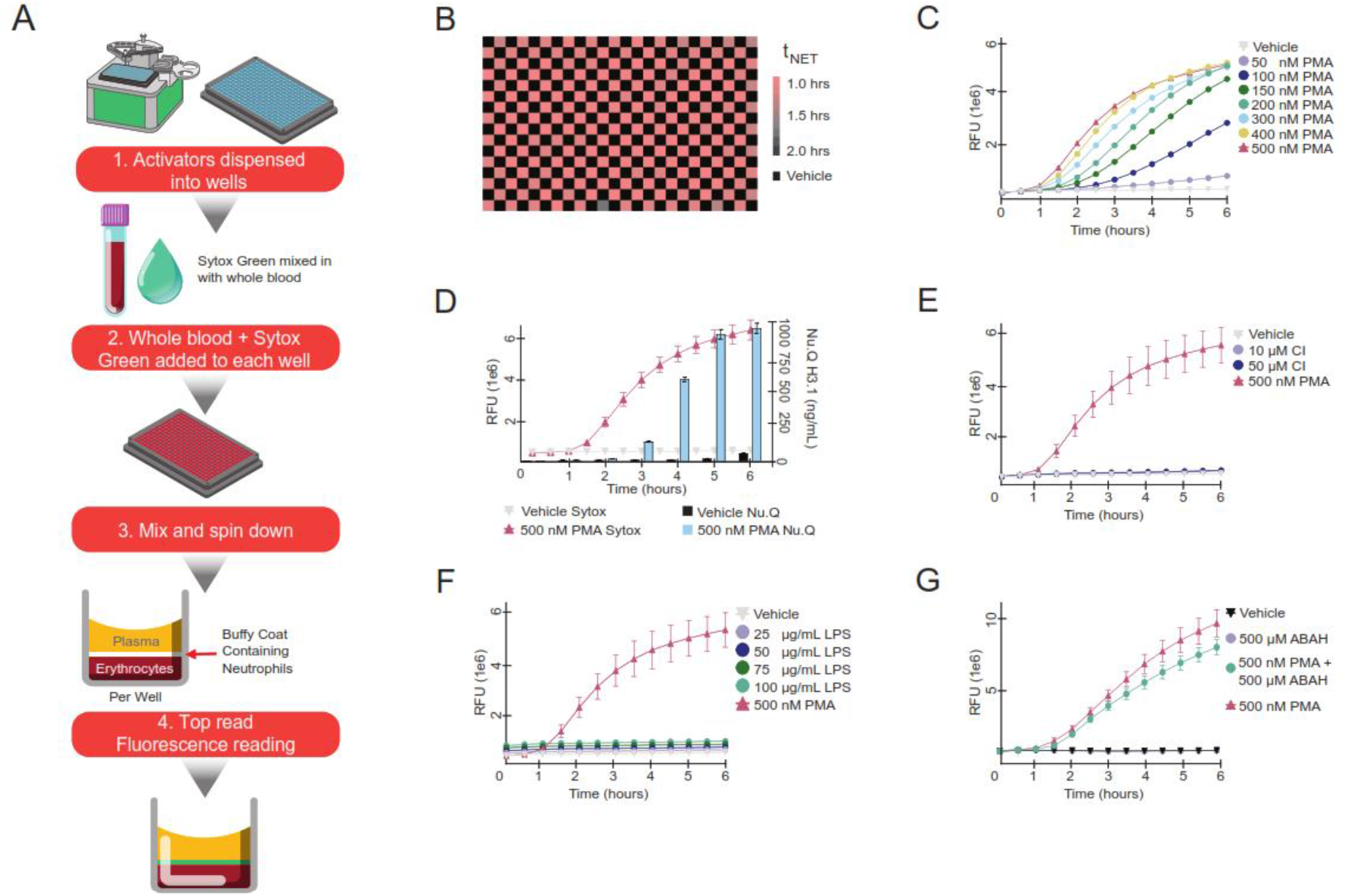
High-throughput ex-vivo NETosis screening method. (A) Workflow for the *ex-vivo* system. Briefly, reagents excluding whole blood are dispensed into a 384-well plate using the MANTIS system (Formulatrix). Whole blood is mixed with Sytox Green and dispensed into the same plate. The plate is mixed and centrifuged for cell separation followed by top read fluorescence. (B) The assay used 500 nM PMA to activate NETosis and the time of NETosis (t_NET_) was measured to illustrate consistency across the plate. (C) PMA titration (·) and a vehicle control (grey ▾) using the *ex-vivo* system was completed over six hours. (D) With blood obtained from one donor, the high-throughput method was compared to H3.1 Nu.Q® where Sytox signal is plotted on the left axis as RFU for the control (grey ▾) and 500 nM PMA (red ▴) and H3.1 Nu.Q® signal on the right axis as ng/mL for control (black bars) and 500 nM PMA (blue bars) at each time point. (E) Time course for 10 or 50 µM CI (light and dark purple ·) compared to 500 nM PMA (red ▴) and a vehicle control (grey ▾) . (F) Time course for 25-100 µg/mL LPS (·), compared to 500 nM PMA (red ▴) and a vehicle control (grey ▾). (G) Time course of NET activation with PMA alone or following pre-incubation with 500 µM of the Myeloperoxidase inhibitor (ABAH) for 45 minutes before the addition of 500 PMA, with ABAH (gray ·) or vehicle (black▾) alone as controls.

To evaluate consistency across the plate and limit potential artifacts (edge effects, temperature variation, and oxygenation), a series of optimizations were performed across an entire 384 plate using 500 nM of PMA to induce NETosis (**Figure 2B**). Experimental parameters were optimized to ensure that small changes in fluorescent intensity at the onset of NETosis could be reliably detected. The time of NETosis (t_NET_), was calculated as the time at which the fluorescent signal exceeded 3.3 × the standard deviation (sd) of a vehicle treated control group. Assay optimization focused on reducing the sd in t_NET_ to +/- 5 minutes across the plate. We then performed a randomized PMA titration across the plate and showed a dose response curve of NETosis induction (**Figure 2C**). Nucleosome release was measured in plasma isolated from treated whole blood in parallel using the H3.1 Nu.Q^®^ NETs assay (Volition) and showed the time course of nucleosome release followed the Sytox green signal (**Figure 2D)**. In this high-throughput, induced NETosis model we found that both CI and LPS failed to induce NETosis (**Figure 2E and Figure 2F**), reflecting the results seen in whole blood (**Figure 1G**). MPO inhibition by ABAH had modest ability to delay the onset of NETosis (**Figure 2G and Supplementary Figure 2A**) consistent with the inhibition of NETosis, as noted in isolated neutrophils. Furthermore, treatment with the ROS and NADPH oxidase inhibitor, Diphenyleneiodonium chloride, (DPI) showed inhibition of DNA release in both Isolated neutrophils and high-throughput system (**Supplementary Figure 1D** and **2B, respectively**).

### Variability of NETosis Profiles

The high-throughput NETosis model was used to screen candidate NETosis regulators selected based on reported association with NETosis or neutrophil biology (**Supplementary Table 1**).

Design of Experiments (DOE) was performed to combine selected factors in an iterative screen to identify the minimal candidate pool required to induce DNA release (**Figure 3A**). Initially, 6-19 factors were combined into individual wells with each factor represented in approximately half of the wells. Four distinct fluorescence patterns were observed following treatment with the various combinations: no response; initial response with early plateau; delayed response with continued gradual increase and initial response with secondary response with continued increase (**Figure 3B**). To assess the relative contribution of each factor to the fluorescence signal at each time point, we down sampled by block averaging into 30-minute intervals, calculated the first derivative, and performed standard least squares multivariate regression modeling (**Figure 3C**). The contribution of each factor across each of the wells was used to determine the potential extent the specific factor contributed to the signal. **Figure 3D** shows examples of four observed patterns: C5a and TNF-α contributed to a rapid onset of DNA release, with TNF-α having a secondary protracted effect on the system. Interleukin-5 (IL-5) did not appear to have a significant impact on fluorescence signal and Interleukin-1β (IL-1β) appeared to reduce the fluorescence signal, indicating potential inhibition or buffering capacity. We performed two rounds of screening in blood from two different healthy donors (n=4) and found that TNF-α, LT-α, IFN-γ, GM-CSF, LTB4, C5a, and Ferritin were predicted to consistently contribute to increased Sytox Green signal across donors whereas LPS and fMLP only contributed in one of the two screens (**Figure 3E**). Molecules with negative or no effect on signal were removed for subsequent NETosis inducer screens. Ferritin was observed to consistently contribute to an increase in signal but was removed from subsequent screens as it was horse derived.

**Figure 3.**
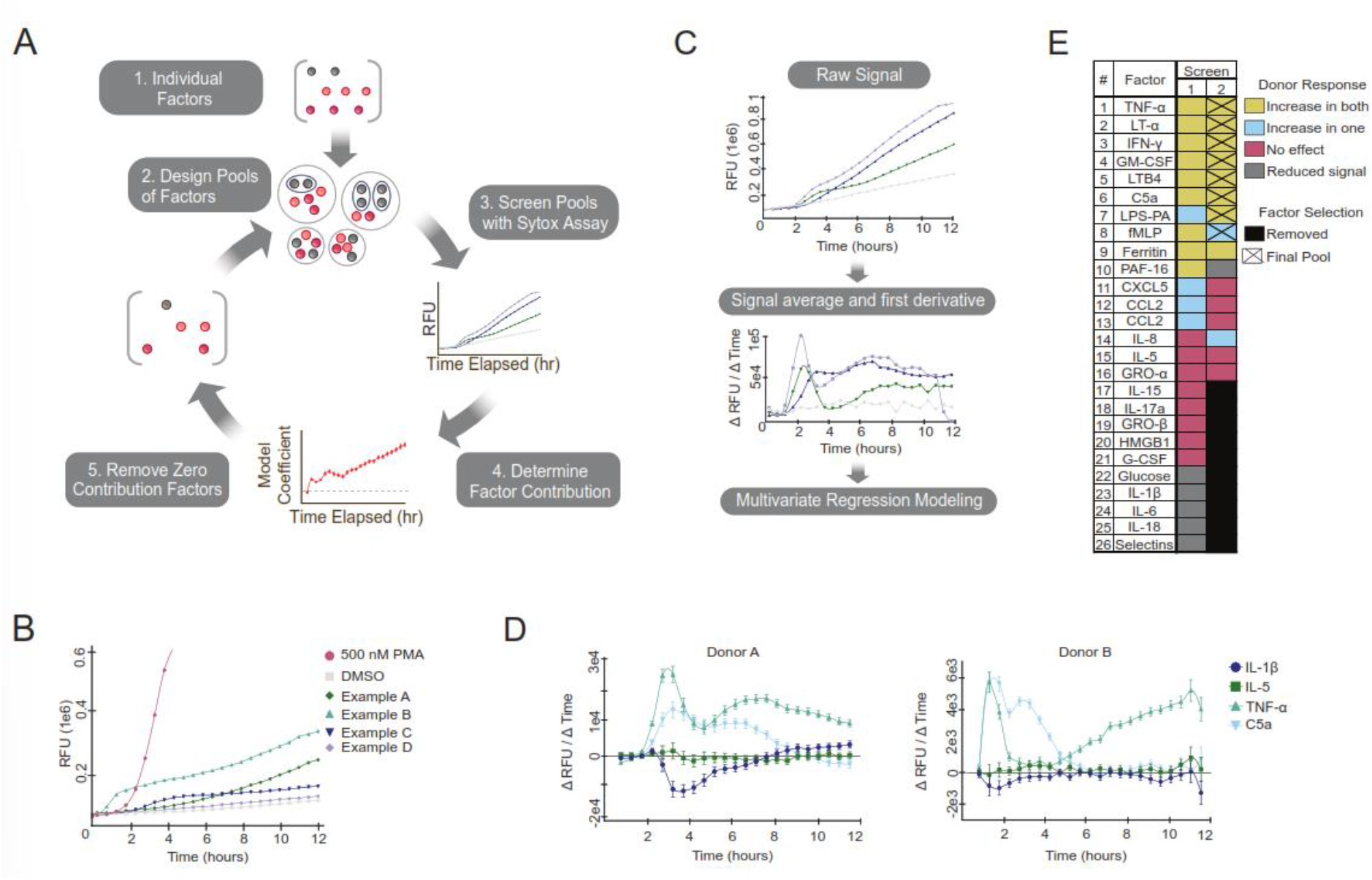
Sequential pooling to identify physiologically relevant NETosis Regulators. (A) Design of the multi-factor screening process. Briefly, individual factors were pooled and tested using the high-throughput screening method and Sytox Green signal was measured. The contribution of each factor was determined and factors which did not increase NETosis were removed, and new pools designed. (B) Four signal profiles were identified for the various pools. The distinct signals were compared to a DMSO vehicle control (◼) and 500 nM PMA (·). Example A (dark green ♦) showed delayed response with continued gradual increase, Example B (▴) showed an initial response with secondary response with continued increase, Example C (▾) showed an initial response with early plateau, and Example D (grey ♦) showed no response. (C) Raw RFU signal was block averaged over 30-minute intervals followed by the first derivative. Afterwards, multivariate regression modeling was completed to determine factor contributions in each pool. (D) An example of the multivariate regression (Figure 3C), highlighting four common factors in 2 donors: C5a (▾) and TNF-α (▴) which showed contribution to the NETosis signal, IL-5 (◼) which had limited response, and IL-1β (·) which showed decreased NETosis signal. (E) The list of factors tested and their responses. Two unique donors were tested in each screen (total n=4) and the final pool was indicated with an X.

### Biological Relevant NETosis Induction Consistency Across Donors

Levels of endogenous cytokines vary across individuals and in response to environmental stimuli contributing to natural variability in innate immune response. Inter-donor variability of NETosis induction was evaluated using the eight selected factors. To normalize the effect each combination of factors had across donors: we performed a Boolean transformation of the first derivative change in RFU over the time course with a threshold cutoff at 3 SD × of the background signal (**Figure 4A**). We tested combinations of the selected 8 factors (TNF-α, LT-α, IFN-g, GM-CSF LTB4, C5a, LPS, and fMLP) in six donors and found signal consistently increased across donors with more factors (**Figure 4B**). Four factors increased NETosis in most of the six donors tested with two distinct patterns: early onset between 30 minutes and two hours, followed by a later increase after three hours. Female donors were generally less responsive than males, especially with earlier onset NETosis (**Figure 4C**). Either TNF-α or LT-α was required for consistent increase in signal across donors. In their absence, the remaining six compounds failed to induce a consistent signal (**Figure 4D**). Interestingly, C5a or LPS appeared to be critical for the early onset signal (30 minutes to 2 hours) (**Figure 4D**). To investigate the relative role of each factor, we selected a pool of five factors (LT-α, GM-CSF, C5a, LPS, and fMLP) which was a pool containing the fewest factors that induced early, but not late, DNA release in all donors tested, hypothesizing the secondary response could reflect mechanisms other than NETosis. The impact of each factor in combination with the other factors across the six donors is shown in **Figure 4E**. These results highlighted differences between male and female donors, for example, it appeared GM-CSF and LPS were less important for the early response in males compared to females and that male donors were more likely to undergo NETosis with less stimulation compared to the female donors with the five-factor pool. TNF-α was also tested in place of LT-α and there was signal in both early and late induction, so it was not used in subsequent pools (**Supplementary Figure 3**).

**Figure 4.**
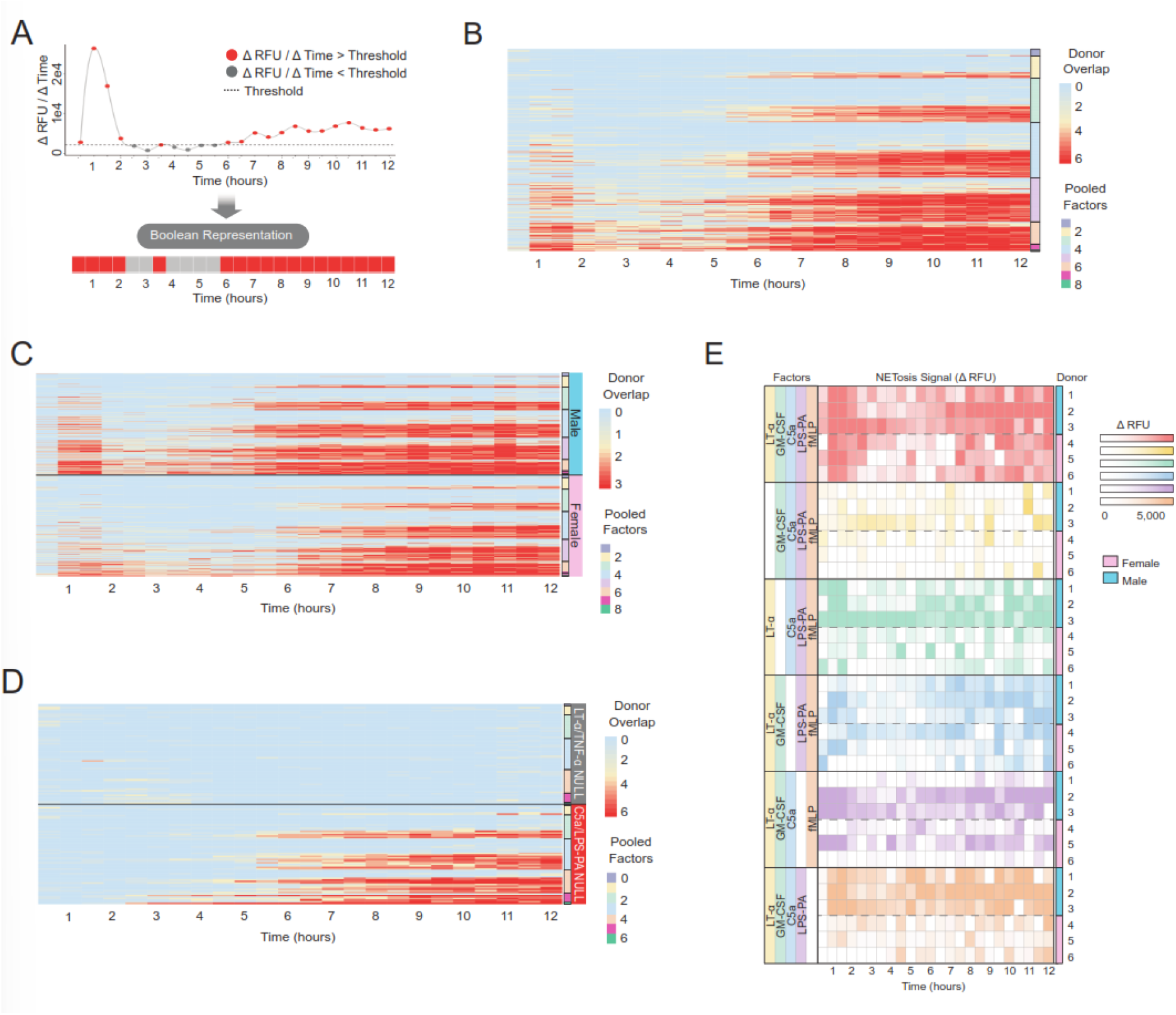
Impact of individual factors within pools across donors. (A) Example of Boolean (binary) representation of the first derivative NETosis signal. If signal was above the threshold (3 × Standard Deviation of Vehicle Signal – dashed line), it was marked as positive (red ·) and converted to a 1, otherwise it was marked as negative (grey ·) and converted to a 0. (B) Boolean representation of NETosis signal over six healthy donors. The factors were pooled in a full factorial manner, such that all possible combinations were present and the number of factors in each pool is indicated. (C) Boolean representation of NETosis signal was separated between male (blue bar) and female (pink bar) donors. (D) Boolean representation of NETosis signal in factor pools when either LT-α and/or TNF-α (top – grey bar) or C5a and/or LPS (bottom – red bar) are not present. (E) Using a limited pool of 5 factors, the change in NETosis signal over background signal is shown upon sequential removal of one factor. Each colored panel reflects a different pool and each row is an individual donor. Red indicates all factors present, yellow removed only LT-α, green removed only GM-CSF, blue removed only C5a, purple removed only LPS, and orange removed only fMLP.

### Minimal factors required for NETosis induction

To determine whether individual donors responded to the same combination of compounds over time we had two donors undergo multiple blood draws over a month period and found general consistency for NETosis initiation with the five factor pool (LT-α, GM-CSF, C5a, LPS, and fMLP) across and within donors, with a pool of LT-α, C5a and fMLP giving the most consistent result with the fewest number of factors (**Figure 5A**). Variability was reduced as more factors were utilized, however, we observed a possible set of three factors (LT-α, C5a, and fMLP) which had consistency and similar results as those pools which included either GM-CSF and or LPS.

**Figure 5.**
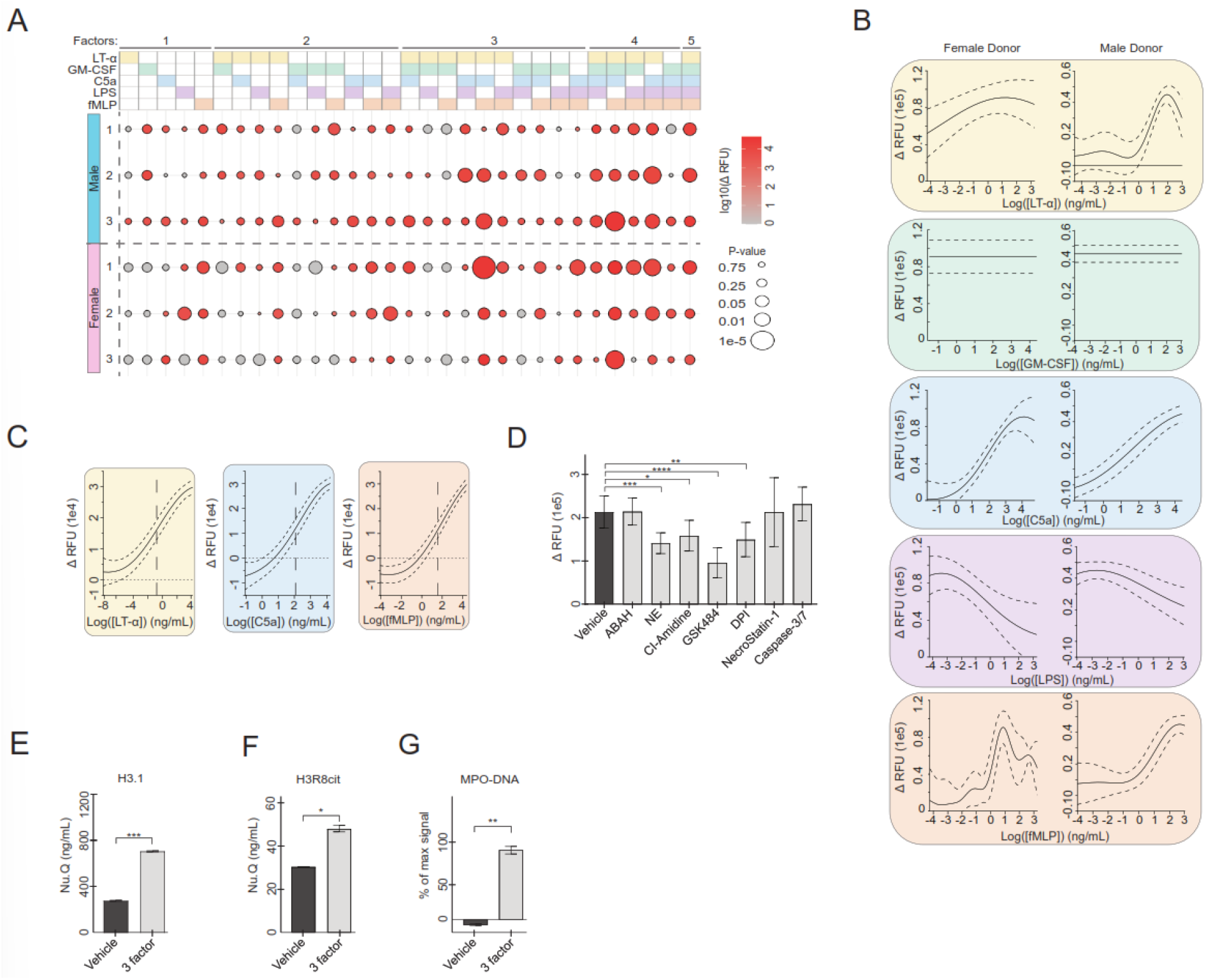
Determination of minimal factors required for NETosis induction in *ex-vivo* model. (A) A male (blue bar – left) and female donor (pink bar – left) were tested at three time points (over 30 days) with a full factorial of the five-factor pool (LT-α, GM-CSF, C5a, LPS, and fMLP). Factors are indicated on the top panel where the filled in color denotes the factor was present in that column (LT-α – yellow, GM-CSF – green, C5a – blue, LPS – purple, and fMLP – orange). Decreasing p-value is indicated by increasing circle size and raw change in the NETosis signal (RFU) after 4 hours is indicated by color intensity. The number refers the draw number (left) for each donor. (B) Gaussian process regression results for combinations of 5-factors at varying concentrations, with dose-response shown for each factor given each other factor is at concentrations giving the maximal response. Delta RFU after 4 hours is shown on the y-axis and log concentration of the factor that is varied (ng/mL) is shown on the x-axis. 95% confidence is shown as dashed lines. (C) EC_50_ concentrations were determined after removing the two factors which showed limited response (GM-CSF and LPS). 95% confidence is shown in the dashed line. The calculated EC_50_ values indicated by a vertical dashed line were: LT-α: 0.3 ng/mL, C5a: 200 ng/mL, and fMLP: 60 ng/mL. (D) Using the pool of three factors at optimized concentrations, inhibitors targeting the following were compared to a Null treatment: 500 µM MPO (ABAH), 100 µM NE (NEi II), 500 µM pan-PAD (CI-Amidine), 100 µM PAD4 (GSK484), 25µM NADPH Oxidase (DPI), 200 µM Receptor-interacting protein kinase 1 (NecroStatin-1), and 750 µM Caspase 3/7 (Caspase-3/7 Inhibitor I). * indicates p < 0.05, ** indicates p < 0.01, *** indicates p < 0.001, and **** indicates p<0.0001. (E-G) Plasma was isolated from whole blood treated with vehicle or a pool of LT-α, C5a, and fMLP and H3.1 Nu.Q® was measured (E), H3R8cit Nu.Q® (F), MPO-DNA represented as the percentage of max OD signal of the assay (G). * indicates p < 0.05, ** indicates p < 0.01, *** indicates p < 0.001.

Due to this, space-filling DOE and Gaussian Process Regression was used to evaluate the concentration dependence of each factor in blood samples from two healthy donors (**Figure 5B**). We found that GM-CSF did not play a significant role in either donor, while LPS appeared to decrease the overall observed signal in a dose-dependent manner (**Figure 5B and Supplementary Figure 4A**). Consequently, we removed GM-CSF and LPS from the pool to generate the minimal combination of factors required to induce an increase in NETosis.

All three remaining factors were required to generate an increase in DNA release associated Sytox Green signal (**Figure 5C**), with the EC_50_ of C5a and fMLP being within 3-4x of published values for neutrophil depolarization and ROS production, respectively, and the EC_50_ for LT-α being lower than what has been reported for ROS production **(Table 1)**. To confirm the signal we were seeing upon treatment with these natural triggers was a result of NETosis, we treated whole blood with the optimized concentrations of LT-α, C5a, and fMLP, along with a variety of inhibitors. We showed a significant decrease in signal when inhibitors targeting Neutrophil Elastase II, peptidylarginine deiminase (PAD), PAD4, and DPI were included (**Figure 5D**).

**Table 1:**
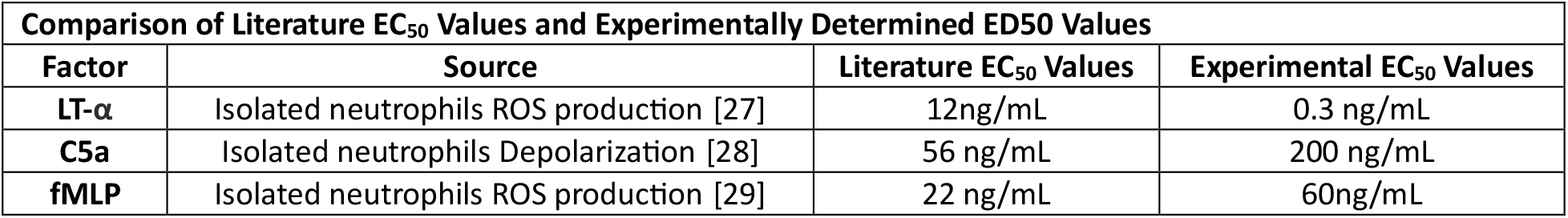
Summary of EC50 values reported in the literature and identified in the current study.

However, there was no change in signal when an apoptosis inhibitor (Caspase 3/7 inhibitor I) or necroptosis inhibitor (necrostatin-1) were included. In addition, we show that following treatment with the three factor pool, there is an increase in H3.1 Nucleosomes (**Figure 5E**), H3R8citrulline (**Figure 5F**), and MPO-DNA (**Figure 5G**) further supporting NETosis induction.

## Conclusion

Physiologically relevant models mimicking endogenous NETosis have the potential to enable mechanistic investigation of the complex signaling underlying NETosis activation. Our representative *ex-vivo* model offers the opportunity for screening therapeutic interventions under disease mimetic conditions which reflect underlying conditions which may predispose patients to adverse outcomes. To our knowledge, we show for the first time the *ex-vivo* induction and real time kinetic read out of NETosis using naturally occurring molecules in a whole blood system. We found that activation with TNF-α or LT-α was required for rapid onset of NETosis, highlighting the importance of these molecules in sepsis and other autoimmune diseases. Indeed, TNF-α has been shown to be elevated in sepsis and is a putative therapeutic target [20].

Interestingly, we found that CI and LPS showed a differential ability to induce NETosis in whole blood compared to isolated neutrophils, which could be due to the presence of an additional cell type or factor present in whole blood, and reflects the complexity of the whole blood model compared to isolated neutrophils. Mol et al, previously reported that treating neutrophils with pairs of factors similar to what we identified (GM-CSF, fMLP, TNF, and LPS), resulted in neutrophils displaying a variety of neutrophil associated behaviors (e.g. ROS production, degranulation, phagocytosis) but they were not able to induce NETosis [15]. Our ability to induce NETosis in our *ex-viv*o model using no less than three different factors, demonstrates the complex signaling needed for NETosis to occur, and suggests that there are components present in whole blood that are important in NETosis regulation. Whole blood has a variety of proteins and other cell types (e.g. platelets and macrophages) which are known to help regulate NETosis and additional work is necessary to determine whether these cells play a role in the NETosis we observe in our *ex-vivo* model.

NETs play a critical role as part of the innate immune response to infection, immobilizing pathogens to prevent dissemination and clearing them from circulation [21]. However, excessive NETosis can be pathogenic and lead to host directed bystander effects and thrombosis [22]. Thus, NETosis can be both beneficial and detrimental [22]. The differential time courses and magnitudes of NETosis seen in our *ex-vivo* whole blood model in response to specific combinations of pro-inflammatory compounds may be correlated to beneficial vs pathogenic NETosis (**Figure 3B**). It is possible that the compound combinations in which the amount of NETosis was minimal represents the beneficial NETosis, whereas conditions that led to higher release may reflect pathogenic NETosis. Furthermore, some combinations trigger a biphasic release pattern with a moderate early minimal response followed by a secondary exacerbated response, which could be reflective of a positive feedback loop in which nucleosomes released from the early NETs further stimulate additional NETosis [23]. DNA and nucleosome release are markers of NETosis but can also measure extracellular traps from other immune cells like eosinophils [24, 25]. We show that MPO and H3R8Cit are expressed following NETosis induction but future studies will address whether different combinations of factors trigger different types of extracellular trap formation or from different cell populations. Differentiating these patterns provide an opportunity for biomarker identification as well as therapeutic intervention for future study.

In our initial screens we found that pools with molecules that interact with similar cell-surface markers/signaling cascades induced consistent responses across donors, which suggested a potential convergence of required signaling pathways. At least three naturally occurring factors in combination (LT-α, fMLP and C5a) were necessary to consistently induce NETosis in our system indicating a potential requirement of the activation of TNF Receptor 1, TNF Receptor 2, or LT-β receptors for NETosis to occur [26]. NETosis induction did not occur in the absence of TNF-α or LT-α, underlying the potentially significant roles these factors play in inflammatory disease, and suggests an underlying master regulatory mechanism, such that certain factors are essential but not individually sufficient to trigger NETosis.

These findings emphasize the importance of expanding the understanding of neutrophil physiology in a biologically relevant context. Physiological triggers of NETosis could be used to better understand NET associated disease pathology, risk factors, and potential therapeutic targets, providing novel strategies for disease intervention and treatment. Our novel *ex-vivo* NETosis approach has the potential to be used for screening drug candidates as inhibitors and inadvertent activators of NETosis and through a comprehensive exploration of human NETosis, we can take significant strides toward mitigating the devastating impact of sepsis on global healthcare.

### Data Sharing

Raw data or additional information will be provided upon emails to the corresponding author.

## Authorship Contributions

K. Zukas, J. Cayford, T. K. Kelly, and M. Eccleston conceived and designed the study, K. Zukas, B. Atteberry and F. Serneo performed the experiments, A. Retter provided clinical guidance and data interpretation. All authors participate in interpretation of data and critical revision of the manuscript for important intellectual content.

Conflict of Interest Disclosures: K. Zukas, F. Serneo, B. Atteberry, J. Cayford and T. K. Kelly are paid employees of Volition. M. Eccleston and A. Retter are paid consultants of Volition.

## Acknowledgments

We would like to thank Alex Hoffman, Scientific Advisory Board Member of Volition, for helpful review of the data and discussion as well as Sarah Erdman for constructive data review and discussions. We would also like to thank Cheryl Kim and Denise Hinz at the La Jolla Institute for Immunology for their FACS sorting work.

---

**Supplemental Table 1:** List of compounds assessed for NETosis regulation activity. All compounds were resuspended according to manufacturer’s recommendation. The per ug cost of compounds were generally a few USD per ug or less, with the exception of LTB4, which was more expensive.

| Compound | Concentration | Catalog # | Source |
| --- | --- | --- | --- |
| Phorbol 12-myristate 13-acetate (PMA) | 50-500nM | P1585 | Sigma-Aldrich |
| Recombinant Human Albumin | 0.1% (w/v) | A9731 | Sigma-Aldrich |
| D-(+)-Glucose | 1mg/mL | G7528 | Sigma-Aldrich |
| Recombinant Human IL-1β | 200ng/mL | RIL1BI | Invitrogen |
| Recombinant Human IL-5 | 200ng/mL | RIL510 | Invitrogen |
| Recombinant Human IL-6 | 200ng/mL | BMS341 | Invitrogen |
| Recombinant Human IL-8 (77aa) | 200ng/mL | PHC0084 | Gibco |
| Recombinant Human IL-15 | 200ng/mL | PHC9154 | Gibco |
| Recombinant Human IL-17a | 200ng/mL | 200-17 | Gibco |
| Recombinant Human IL-18 | 200ng/mL | 10119-HNCE | Sino Biological |
| Recombinant Human TNF-α | 200ng/mL | RTNFAI | Invitrogen |
| Recombinant Human LT-α | 200ng/mL | 300-01B | Gibco |
| Recombinant Human IFN-γ | 1000ng/mL | RIFNG100 | Invitrogen |
| Recombinant Human E-Selectin | 200ng/mL | SRP3034 | Sigma-Aldrich |
| Recombinant Human P-Selectin | 200ng/mL | 561306 | Sigma-Aldrich |
| PAF-16 | 200ng/mL | 511075 | Sigma-Aldrich |
| Recombinant Human GRO-α | 200ng/mL | 300-11 | Gibco |
| Recombinant Human GRO-β | 200ng/mL | PHC1074 | Gibco |
| LTB<sub>4</sub> | 100ng/mL | 23-075-OU | Tocris Bioscience |
| Recombinant Human CXCL5 | 200ng/mL | 300-22B | Gibco |
| Recombinant Human CCL2 | 200ng/mL | PHC1015 | Gibco |
| Recombinant Human CCL3 | 200ng/mL | RP-10923 | Invitrogen |
| Recombinant Human G-CSF | 200ng/mL | PHC2035 | Gibco |
| Recombinant Human GM-CSF | 200ng/mL | RGMCSF20 | Invitrogen |
| fMLP | 400ng/mL | 47729 | Sigma-Aldrich |
| Ferritin (Equine) | 10µg/mL | F4503 | Sigma-Aldrich |
| Recombinant Human HMGB1 | 200ng/mL | ab167718 | Abcam |
| Recombinant Human C5a | 200ng/mL | 300-70 | Gibco |
| LPS (*Pseudomonas aeruginosa*) | 1µg/mL | L8643 | Sigma-Aldrich |

