## Supplemental Figures and legends for "Rapid High-Throughput Method for Investigating Physiological Regulation of Neutrophil Extracellular Trap Formation"

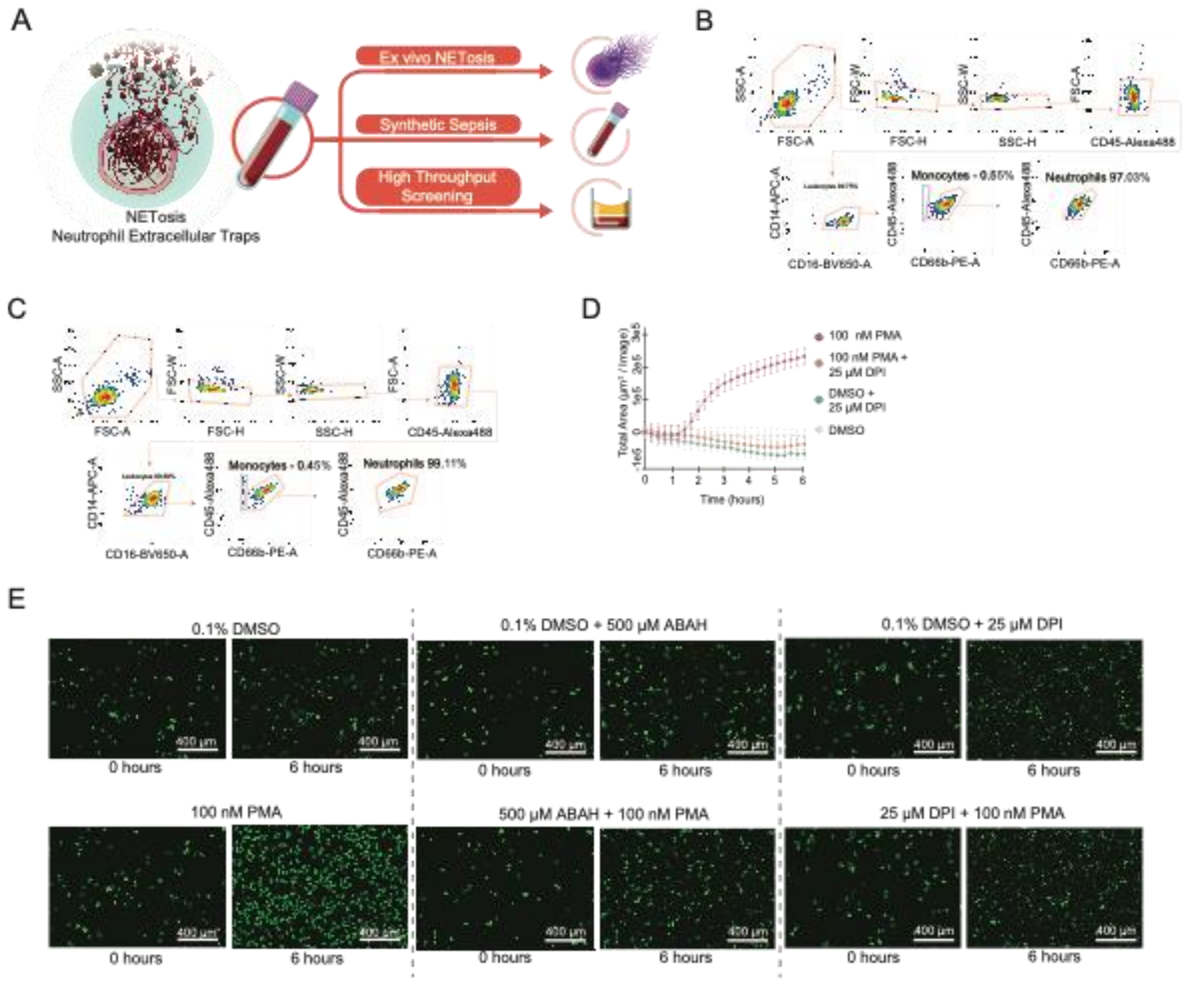

**Supplementary Figure 1:**
(A) NETosis induction overview. (B-C) Isolated neutrophils were gated through FSC-A vs SSC-A and then further by FSC-H vs FSC-W and SSC-H vs SSC-W. CD45+ cells were selected and further separated to obtain CD16+CD14+ leukocytes. To remove monocytes (CD45+CD66b-), CD45+CD66b+ was used to confirm high purification of neutrophils ~97% in donor 1 and ~99% in donor 2. (D) 25 µM diphenyleneiodonium chloride (DPI) inhibition on primary isolated neutrophils using Cytotox green dye on an S3 Incucyte imaging system following induction with 100 nM PMA (red ·), 25 µM DPI + 100 nM PMA (orange ·) , 25 µM DPI + DMSO (green ·), and a DMSO vehicle control (grey ·) on primary isolated neutrophils. (E) Representative images from the Incucyte taken at the start of the assay (left) and after six hours (right) for Figures 1B, 1C, and Supplement Figure 2D.

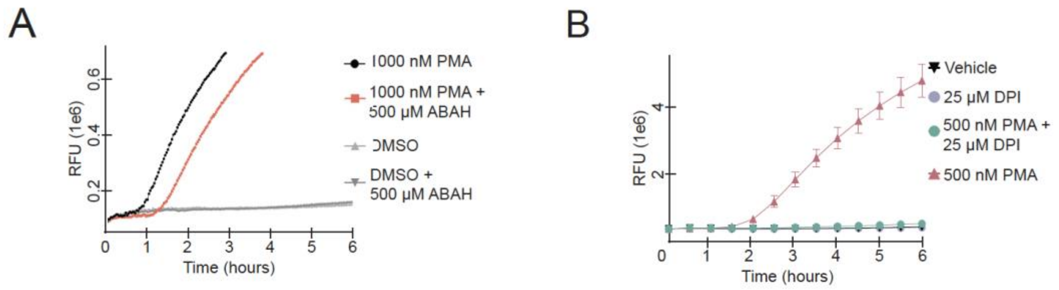


**Supplementary Figure 2:** (A) Using the *ex-vivo* system and high sensitivity instrument gain to detect early changes in fluorescence and 1000 nM PMA (black ·) compared to treatment with 1000 nM PMA with 500 µM ABAH (red ■). Also included was a DMSO vehicle control ( ▲ ) and a DMSO vehicle control with 500 µM ABAH ( ▼ ). The whole blood was incubated at room temperature for 45 minutes with or without the ABAH added prior to addition of PMA. PMA associated signal in saturated after multiple hours and no longer plotted on graph. (B) Using the gain settings as in Figure 2 Sytox green signal following treatment with 500 nM PMA (red ▲ ) compared to 500 nM PMA with 25 µM diphenyleneiodonium chloride (DPI) (green ·). Also included was a DMSO vehicle control (black ▼ ) and a DMSO vehicle control with 500 µM ABAH (grey ·). The whole blood was incubated for 45 minutes with or without the ABAH added prior to addition of PMA.

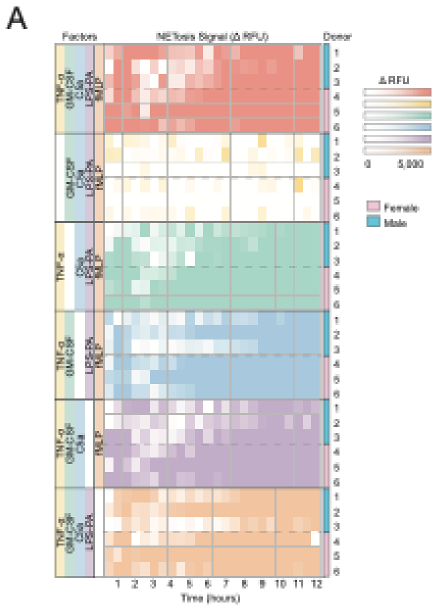


**Supplementary Figure 3** (A) Similar to Figure 4E, but LT-α was replaced with TNF-α. Using a limited pool of 5 factors (left panel), the change in NETosis signal over background signal is shown upon sequential removal of one factor. Each colored panel on the right reflects a different pool and each row is an individual donor. Red indicates all factors present, yellow removed only TNF-α, green removed only GM-CSF, blue removed only C5a, purple removed only LPS, and orange removed only fMLP.

A

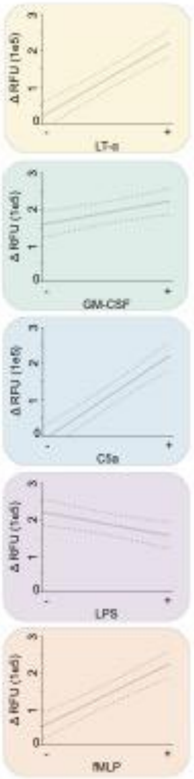


**Supplementary Figure 4** (A) Standard least squares regression analysis of full-factorial donor data (Figure 5A), assigning the donor and day as nested random variables and each of the five factors (LT-α, GM-CSF, C5a, LPS, and fMLP) as independent variables. The effect of each factor is shown given that each factor absent or present in a combination that gives the optimal signal. Y-axis indicates the difference in signal intensity after 4 hours using the high-throughput method. X-axis indicates whether the factor was present or not ("-" indicates removal of the factor and "+" indicates the factor was present). Dashed lines indicate 95% confidence intervals.
